# Development of attention networks from childhood to young adulthood: A study of performance, intraindividual variability and cortical thickness

**DOI:** 10.1101/2020.10.05.326835

**Authors:** Rune Boen, Lia Ferschmann, Nandita Vijayakumar, Knut Overbye, Anders M. Fjell, Thomas Espeseth, Christian K. Tamnes

## Abstract

Human cognitive development is manifold, with different functions developing at different speeds at different ages. Attention is an important domain of this cognitive development, and involves distinct developmental trajectories in separate functions, including conflict processing, selection of sensory input and alertness. In children, several studies using the Attention Network Test (ANT) have investigated the development of three attentional networks that carry out the functions of executive control, orienting and alerting. There is, however, a lack of studies on the development of these attentional components across adolescence, limiting our understanding of their protracted development. To fill this knowledge gap, we performed a mixed cross-sectional and longitudinal study using mixed methods to examine the development of the attentional components and their intraindividual variability from late childhood to young adulthood (n = 287, n observations = 408, age range = 8.5–26.7 years, mean follow up interval = 4.4 years). The results indicated that executive control stabilized during late adolescence, while orienting and alerting continued to develop into young adulthood. In addition, a continuous development into young adulthood was observed for the intraindividual variability measures of orienting and alerting. In a subsample with available magnetic resonance imaging (MRI) data (n =169, n observations = 281), higher alerting scores were associated with thicker cortices within a right prefrontal cortical region and greater age-related cortical thinning in left rolandic operculum, while higher orienting scores were associated with greater age-related cortical thinning in frontal and parietal regions. Finally, increased consistency of orienting performance was associated with thinner cortex in prefrontal regions and reduced age-related thinning in frontal regions.

## Introduction

Three decades ago, Posner and Petersen (1990) proposed to divide attention into subsystems that carry out separable attentional functions. This theoretical framework was later used to develop the ANT (Fan et al., 2002), which combines the Eriksen flanker task (Eriksen & Eriksen, 1974) and the cued reaction time task (Posner, 1980) into a single behavioral paradigm. In the ANT, executive control is defined as resolving conflict among responses; orienting as the selection of information from sensory input; and alerting as achieving and maintaining an alert state (Fan et al., 2002, 2009). The efficacy of the attentional networks that carry out these functions during the ANT have been investigated by comparing the reaction times in the incongruent trials (executive control), spatial cue trials (orienting) and double cue trials (alerting) to the reaction times in congruent trials, center cue trials and no cue trials, respectively (Fan et al., 2002). Here, the efficacy of the executive control network is estimated by how fast the participant can resolve and react to the conflict induced by incongruent flankers, where lower executive control scores are indicative of better conflict resolution abilities. The efficacy of the orienting network is estimated by the reduced reaction time following spatial cues, where higher orienting scores indicate better ability to utilize spatial information to orient and react to the correct target location. The efficacy of the alerting network is estimated by the reduced reaction time following alerting cues, where higher alerting scores indicate higher vigilance before the appearance of a target stimuli or, alternatively, difficulty of maintaining alertness when there is no cue.

The ANT has been used to investigate age-related differences in attention in different parts of life (e.g. Mahoney et al., 2010; Rueda et al., 2004), as well as attentional functions in clinical groups, including individuals with ADHD (Adólfsdóttir et al., 2008; Mogg et al., 2015), schizophrenia (Spagna et al., 2015), and autism (Fan et al., 2012). Although there have been several developmental studies on the attentional components in middle and late childhood (Federico et al., 2017; Mezzacappa, 2004; Mullane et al., 2016; Pozuelos et al., 2014; Rueda et al., 2004), there is a limited number of longitudinal studies directly investigating developmental changes (but see Lewis et al., 2016; Suades-González et al., 2017). Thus, the late developmental trajectories of these attentional components remain poorly understood. Finally, while we know that the brain undergoes substantial structural changes across adolescence (Dennis & Thompson, 2013; Tamnes & Mills, 2020), we have limited understanding of how the development of specific attentional components relate to these structural changes.

Previous research suggests that the attentional components have different developmental trajectories during childhood (Federico et al., 2017; Lewis et al., 2016; Mezzacappa, 2004; Mullane et al., 2016; Pozuelos et al., 2014; Rueda et al., 2004). Executive control seems to develop until early adolescence (Baijal et al., 2011; Mullane et al., 2016; Rueda et al., 2004), possibly beginning to stabilize around late adolescence (Waszak et al., 2010), orienting seems to be stable across middle childhood (Lewis et al., 2016; Mullane et al., 2016; Rueda et al., 2004; Suades-González et al., 2017), while alerting has been shown to continue to develop from early childhood into late childhood (Lewis et al., 2016; Mullane et al., 2016; Pozuelos et al., 2014). However, it should be noted that the alerting scores typically decrease with age, possibly because younger children do poorly in no cue trials (Rueda et al., 2004). Moreover, the abovementioned studies have typically focused on overall reaction time and accuracy measures, but not intraindividual variability; i.e. variability across trials or sessions of the same task (MacDonald et al., 2009; Williams et al., 2005). Previous research has shown large age effects for intraindividual variability in both congruent and incongruent trials of the flanker task (Tamnes et al., 2012) and for choice reaction time and simple reaction time tasks during development (Dykiert, Der, Starr, & Deary, 2012), documenting improved consistency with increasing age. By using the different trial types in the ANT, it is possible to investigate the development of the intraindividual variability in each of the three attentional components, henceforth referred to as executive control variability, orienting variability and alerting variability. Similar to the reaction times during the ANT, it seems plausible that intraindividual variability in incongruent trials, spatial cue trials and double cue trials will show a continuous reduction with age, suggesting that a decrease in executive control variability and an increase in orienting variability and alerting variability are likely to be observed with increasing age. This field of research is important as within-task intraindividual performance has been shown to provide insight into temporal fluctuations of cognitive functioning (Bielak et al., 2010; MacDonald et al., 2006), and is found to be larger among children with attentional problems compared to controls (Epstein et al., 2011).

In parallel to the development of the attentional networks, the brain also undergoes substantial structural changes (Fjell et al., 2015; Herting et al., 2018; Tamnes et al., 2017; Vijayakumar et al., 2016; Wierenga et al., 2014), but studies linking the two are lacking. Previous functional MRI and electroencephalography (EEG) studies have investigated ANT associated brain activity among both adults (Fan et al., 2005, Galvao-Carmona et al., 2014; Neuhaus et al., 2010) and children (Konrad et al., 2005, Gopalan et al., 2019), and have reported that the different attentional networks recruits different brain regions. Executive control has been associated with activity in the anterior cingulate and bilateral prefrontal cortical regions (Fan et al., 2005; Konrad et al., 2005), orienting with activity in bilateral superior parietal cortical regions, and alerting with activity in thalamic, inferior parietal and prefrontal cortical regions (Fan et al., 2005). Structural MRI studies are scarce, but a cross-sectional study by Westlye et al (2011), which included 286 adults 20-84 years of age, showed a positive association between executive control and cortical thickness of the anterior cingulate and inferior frontal cortical regions. They also reported negative associations between alerting and thickness in superior parietal cortical regions, while no relations between orienting and cortical thickness were found (Westlye et al., 2011). While this study provided insight into the structural correlates of the attentional networks in adults, the associations between the development of these attentional networks and structural brain maturation remain unknown.

The present study aimed to examine the 1) developmental trajectory, 2) intraindividual variability, and 3) structural cortical correlates of executive control, orienting and alerting from late childhood to young adulthood. Based on previous behavioral studies, we expect executive control performance improvements to decelerate (i.e. initial decrease that slows down) with age (Baijal et al., 2011; Mullane et al., 2016; Waszak et al., 2010), while we expect orienting to remain stable from late childhood to young adulthood (Rueda et al., 2004; Suades-González et al., 2017). Previous results have shown a decrease in alerting during childhood (Lewis et al., 2016; Pozuelos et al., 2014), possibly driven by poor performance in no cue trials among the youngest participants (Rueda et al., 2004). However, as alerting scores may reflect either difficulty in maintaining an alert state in cue trials or efficient use of alerting cues (Posner, 2008), and we do not know how these processes change across adolescence, no prediction about alerting could be made within the current age range. Furthermore, as intraindividual variability shows large age-related effects across childhood to young adulthood (Dykiert et al., 2012; Tamnes et al., 2012), we also expect a continuous development of the intraindividual variability measures of the attentional components across adolescence. Since we expect increased consistency in incongruent trials, spatial cue trials and double cue trials with age, we predict that the executive control variability scores will decrease with age and that orienting variability will increase with age. However, if the previous notion that the alerting effect is highly influenced by the performance in no cue trials is true, it seems likely that this will be reflected in the variability scores as well, hence no prediction about the direction could be made. Finally, we hypothesize that the attentional components and their intraindividual variability to be related to cortical thickness and age-related cortical thinning in the ANT associated brain regions (Fan et al., 2005; Konrad et al., 2005; Westlye et al., 2011). Thus, we expect that thinner cortex and faster age-related cortical thinning in the anterior cingulate and lateral prefrontal regions to be associated with lower executive control and executive control variability scores. Furthermore, we predict that thinner cortex and faster age-related cortical thinning in the superior parietal region to be associated with higher orienting and orienting variability scores, and that cortical thickness and age-related thinning in parietal and prefrontal regions will be associated with alerting and alerting variability scores.

## Materials and Methods

### Participants

The sample was drawn from the three-wave accelerated longitudinal research project *Neurocognitive Development* (Ferschmann et al., 2019; Tamnes et al., 2010, 2013). The study was approved by the Regional Committee for Medical and Health Research Ethics. Initially, children and adolescents were recruited through newspaper ads and local schools, while new participants were recruited through social media platforms in the second and third waves to compensate for attrition and to increase the sample size and to account for practice effects. Written informed consent was obtained from all participants over 12 years of age and from a parent of participants under 16 years of age. Oral informed assent was given by children under 12 years of age. At each wave, participants aged 16 years or older and a parent of participants under 16 years were screened with standardized health interviews to ascertain eligibility. Participants were required to be right handed, fluent Norwegian speakers, have normal or corrected to normal vision and hearing, not have history of injury or disease known to affect central nervous system (CNS) function, including neurological or mental disorders and serious head trauma, not be under treatment for a mental disorder, not use psychoactive drugs known to affect CNS functioning, not have had a premature birth (< 37 weeks), and not have MRI contraindications. All MRI scans were evaluated by a neuroradiologist and required to be deemed free of serious injuries or conditions.

A total of 287 participants (153 females) in the age-range 8.5–26.7 years (M = 17.6, SD = 4.1, across all observations) were included in the current study and had ANT data from one or two time points (ANT was not included in the test protocol in the first wave of the *Neurocognitive Development* project). Of these, 166 participants (92 females) had data from one time-point and 121 participants (61 females) had data from two time-points, yielding a total of 408 observations. For participants with data from two time-points, the mean interval between the observations was 4.4 years (SD= 0.4, range= 3.9–5.5). The length of the interval was not related to age (r=.08, p=.36).

### MRI subsample

A subsample of 169 participants (83 females) had quality controlled MRI data (see below for details) from the same scanner and were included for MRI analyses, where 112 participants (60 females) had MRI data from two time points, yielding a total of 281 observations with valid MRI and ANT data. Of the full sample, 117 participants recruited in the third wave were not scanned on the same scanner as the rest of the sample due to transition to a new MRI scanner, and one participant had missing MRI data, thus 118 participants were not included in the MRI analyses.

### The Attention Network Test

We administered an adult version of the ANT (Fan et al., 2002), which consisted of a practice block with 24 trials and three blocks with 96 trials each, for a total of 288 test trials. During assessment, participants were seated in a comfortable chair at approximately 60 cm distance from a 19-inch monitor. For each trial, participants were required to press a key indicating whether a target arrow was pointing to the left or right. The arrow was presented either above or below a centrally located fixation cross. The target arrow was flanked by one of three different types of stimuli: 1) pairs of congruent arrows, 2) pairs of incongruent arrows, or 3) pairs of neutral lines. Each type of flanker stimuli was presented 32 times per block. Furthermore, each trial was preceded by one of four cue conditions, with each variant occurring 24 times in each block: 1) no cue, 2) center cue, 3) double cue, or 4) spatial cue. The cues, when presented, were single (center and spatial cue) or double asterisks replacing (center cue) or accompanying the fixation cross. The size of the fixation cross was approximately 0.5 × 0.5 cm (~0.5°), and the diameter of the asterisks used for cueing was about 0.3 cm (~0.3°). Target arrows were horizontally centered 1.3 cm (~1.2°) below or above the fixation cross. Each trial was initiated by the fixation cross for a random duration of 400-1600 ms. This was followed by the cue stimuli for 100 ms, the fixation cross for 400 ms and then the target stimuli, which remained visible on the screen until response or for 1700 ms. The participants were instructed to focus on both speed and accuracy throughout the session, and the length of the breaks between blocks were controlled by the participant. The experimental procedure was administered using E-prime software (Schneider et al., 2002), and responses were obtained on a PST Serial Response Box. The ANT adult version was chosen instead of the ANT child version (Rueda et al., 2004), as pilot testing suggested that it was more suitable across the current age-range and since it includes more trials per time.

### MRI acquisition

The MRI data was collected at Rikshospitalet, Oslo University Hospital with the same 12-channel head coil on the same 1.5 T Siemens Avanto scanner (Siemens Medical Solutions, Erlangen, Germany) at both time-points. The sequence used for morphological analyses was a 3D T1-weighted magnetization prepared rapid gradient echo (MPRAGE) with the following parameters: repetition time/echo time/time to inversion/flip angle = 2400 ms/3.61 ms/1000 ms/8°, matrix = 192 × 192 × 160, sagittally acquired, field of view = 240 mm, bandwidth = 180 Hz/pixel, voxel size 1.25 × 1.25 × 1.2 mm. Acquisition time of this sequence was 7 minutes and 42 seconds.

### MRI processing

The MRI data underwent whole brain segmentation and cortical surface reconstruction and longitudinal processing in Freesurfer 6.0 (Dale et al., 1999; Fischl et al., 1999, 2002, 2012; Reuter et al., 2012). Cortical thickness was estimated based on the shortest distance between the gray matter/white matter boundary and the cortical surface. This was conducted at each vertex across the cortical mantle. Before statistical analyses, cortical thickness was resampled to 81,924 vertices and the thickness maps were smoothed with a Guassian kernel with a full-width of half-maximum of 10 mm. All MR images were manually inspected by a trained individual and a re-scan from the same session was used if available and if the images were deemed to be of insufficient quality (see Ferschmann et al., 2019 for details)

### Statistical analyses

#### ANT components

To calculate ANT components, we used a ratio procedure with median reaction times based on correct trials. The ratio procedure was chosen to isolate the attention scores from the expected decrease in reaction times with age. We computed the following attention network scores (RT represents the reaction time):

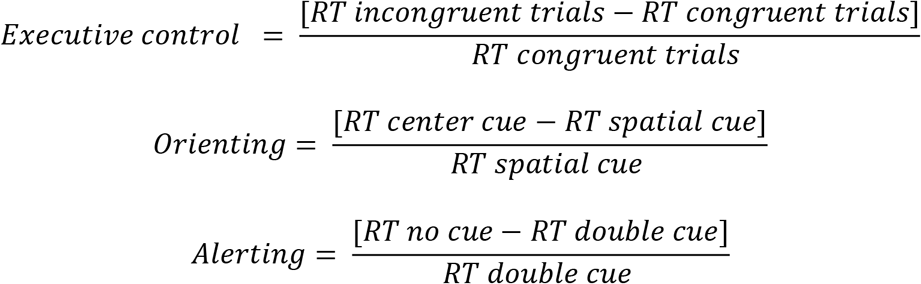

Additionally, we utilized the same ratio procedure, replacing median reaction time with the standard deviation of the median reaction time to obtain a measure of intraindividual variability for each of the ANT components. We computed the following intraindividual variability scores (SD represents the standard deviation):

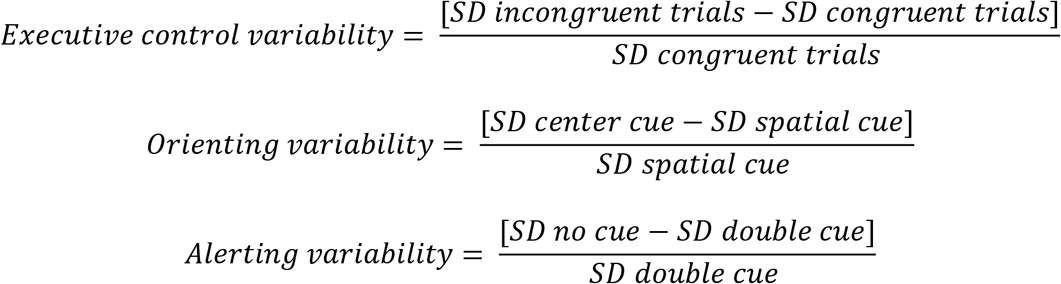

#### Development of the ANT components and their intraindividual variability

We used the nlme package (Pinheiro, Bates, DebRoy, Sarkar, & R Core Team, 2018) in R (R Core Team, 2018) to investigate the development of the attentional components and their intraindividual variability. All the continuous predictors in the general linear mixed models were centered around their mean prior to the analyses. First, we assessed the best fitting developmental model by comparing the following models:

1) null model (Y = intercept + random (participant ID) + error)
2) linear age model (Y = intercept + random (participant ID) + age + error)
3) quadratic age model (Y = intercept + random (participant ID) + age + age^2^+ error)

Second, if one of the age models was a significantly better fit than the null model, we examined whether the best fitting age model was further improved by including sex in the model (e.g., for linear model):

4) Main sex model (Y = intercept + random (participant ID) + age + sex + error)
5) Interaction age^x^sex model (Y = intercept + random (participant ID) + age + sex + age*sex + error)

Models were fitted with maximum likelihood (ML) estimation method. To avoid overfitting, model selection was guided by *p*-values generated by the likelihood ratio (LRT) test, but the Akaike Information Criterion (AIC; Akaike, 1974) was emphasized. More specifically, the more complex model was chosen if the LRT *p*-value dropped below .05 and the AIC value decreased by at least 2. Since starting and ending points of samples can influence the age-related trajectory when using parametric models (Fjell et al., 2010), we used generalized additive mixed models (GAMM) with a cubic spline basis (Wood, 2017) for visualization of the results. The GAMM plots were produced using the following model in R: Y ~ s(age, bs = ‘cr’), where “s” is the function used to define smooth terms, whereby the smooth is a function of “age”, “bs” is the smoothing basis to use, and the “cr” refers to a cubic regression spline. Note that the GAMM plots are used for visualizations only, while the plotted general linear mixed models can be found in the supplementary materials (Supplementary Figures 1 and 2).

#### ANT components and cortical thickness

We conducted vertex-wise analyses in SurfStat (http://www.math.mcgill.ca/keith/surfstat/) to investigate the relationship between behavioral performance in the ANT and cortical thickness. At each vertex, we computed a linear mixed model to investigate the effect of the attentional components and their intraindividual variability scores (ANT_component_), interaction effect of ANT_component_ *age and ANT_component_ *age^2^ using the following model: Y_thickness_ = intercept + ANT_component_ + Sex + Age + Age^2 + ANT_component_ *Age + ANT_component_ *Age^2 + random(participant) + error in SurfStat All the continuous predictors were centered around their mean before running the analyses. To account for multiple comparisons, we used a random field theory correction of p < .05 and a cluster-defining threshold of p < .005. Here, we will report the significant clusters of the main effect of ANT_component_, interactions between ANT_component_ and age, and ANT_component_ and age^2^.

## Results

### Overall behavioral performance

Behavioral data for each trial type are presented in Table 1. The results showed that the participants had > 91% accuracy across all trial types, indicating that the participants understood the task. The mean attentional component scores are presented in Table 2. Across all observations, the descriptive data showed lower RTs in the congruent, spatial cue and double cue compared to the incongruent, center cue and no cue condition, respectively. The positive values for the attentional components indicate that the sample showed the expected effect of the task conditions.

**Table 1.** Accuracy and reaction times for the ANT trial types.

|  | Cue condition |  |  |  | Flanker condition |  |  |
| --- | --- | --- | --- | --- | --- | --- | --- |
|  | Center | Double | No | Spatial | Congruent | Incongruent | Neutral |
| Accuracy | 95.43 | 95.57 | 97.00 | 97.24 | 99.05 | 91.29 | 98.69 |
| (SD) | (4.67) | (4.45) | (3.29) | (3.85) | (1.64) | (8.23) | (1.96) |
| Mean | 485.17 | 474.72 | 519.46 | 437.77 | 452.00 | 554.05 | 452.19 |
| (SD) | (81.27) | (76.14) | (82.97) | (74.78) | (72.78) | (94.33) | (70.06) |

Note: Accuracy is presented in percentage. Mean reaction times are based on median reaction time on correct trials. Accuracy and reaction times are estimated across all observations. SD = standard deviation.

**Table 2.** ANT performance for the sample as a whole, and females and males separately

|  | Total sample |  |  | Females | Males |
| --- | --- | --- | --- | --- | --- |
|  | Mean (SD) | Min | Max | Mean (SD) | Mean (SD) |
| Executive |  |  |  |  |  |
| control | .23 (.07) | .07 | .56 | .23 (.07) | .22 (.07) |
| Orienting | .11 (.05) | -.03 | .26 | .12 (.05) | .10 (.05) |
| Alerting | .10 (.06) | -.06 | .34 | .10 (.06) | .09 (.05) |

Note: SD = standard deviation.

### Development of the ANT components

For executive control, a quadratic age model, with no main effect of sex or age-sex interaction, was found to best fit the data (Table 3). For orienting, the best fitting model was a positive linear model with sex included as a main effect (Table 4), while alerting was best fitted by a negative linear model with no main effect of sex or age-sex interaction (Table 5). The results from the mixed model analyses showed that the three attention networks had three different age models that best fitted the data (Table 6). The GAMM visualizations (Figure 1) indicate a decelerating negative effect of age on executive control, stabilizing around late adolescence, while the GAMM for orienting shows a linear increase across the sample range, with females having higher orienting scores than males. The GAMM slope for alerting, however, indicates a cubic pattern, with a decelerating negative effect of age during adolescence.

**Table 3.** Comparison of models predicting executive control

| Model | Model number | df | AIC | BIC | LogLik | Test | L.Ratio | p |
| --- | --- | --- | --- | --- | --- | --- | --- | --- |
| Null model | 1 | 3 | -1050.57 | -1038.53 | 528.28 | - | - | - |
| Linear age model | 2 | 4 | -1055.78 | -1039.74 | 531.89 | 1 vs 2 | 7.21 | **0.007** |
| Quadratic age model | 3 | 5 | -1059.47 | -1039.41 | 534.74 | 2 vs 3 | 5.69 | **0.017** |
| Main sex model | 4 | 6 | -1058.29 | -1034.22 | 535.14 | 3 vs 4 | 0.82 | 0.366 |
| Interaction Sex*Age<sup>2</sup> model | 5 | 8 | -1057.99 | -1025.90 | 536.99 | 3 vs 5 | 4.52 | 0.211 |

**Table 4.** Comparison of models predicting orienting

| **Model** | **Model**<br>**number** | **df** | **AIC** | **BIC** | **LogLik** | **Test** | **L.Ratio** | **p** |
| --- | --- | --- | --- | --- | --- | --- | --- | --- |
| Null model | 1 | 3 | -1293.55 | -1281.52 | 649.78 | - | - | - |
| Linear age<br>model | 2 | 4 | -1300.10 | -1284.05 | 654.05 | 1 vs 2 | 8.55 | **0.003** |
| Quadratic<br>age model | 3 | 5 | -1299.60 | -1279.55 | 654.80 | 2 vs 3 | 1.50 | 0.220 |
| Main sex<br>model | 4 | 5 | -1309.51 | -1289.46 | 659.76 | 2 vs 4 | 11.42 | **0.001** |
| Interaction<br>sex*age<br>model | 5 | 6 | -1307.52 | -1283.45 | 659.76 | 2 vs 5 | 11.42 | **0.003** |
| Interaction<br>sex*age<br>model | 5 | 6 | -1307.52 | -1283.45 | 659.76 | 4 vs 5 | 0.00 | 0.957 |

**Table 5.** Comparison of models predicting alerting

| Model | Model number | df | AIC | BIC | LogLik | Test | L.Ratio | p |
| --- | --- | --- | --- | --- | --- | --- | --- | --- |
| Null model | 1 | 3 | -1198.03 | -1185.99 | 602.01 | - | - | - |
| Linear age | 2 | 4 | -1204.53 | -1188.49 | 606.27 | 1 vs 2 | 8.50 | **0.004** |
| Quadratic age model | 3 | 5 | -1203.38 | -1183.32 | 606.69 | 2 vs 3 | 0.85 | 0.357 |
| Main sex model | 4 | 5 | -1203.75 | -1183.70 | 606.88 | 2 vs 4 | 1.22 | 0.269 |
| Interaction sex*age model | 5 | 6 | -1202.76 | -1178.69 | 607.38 | 2 vs 5 | 2.23 | 0.328 |

**Table 6.** Model parameters for the best fitting model of the attentional components

| Attention network | Value | S.E | CI (95%)<br>[Lower, Upper] | t-value | p-value |
| --- | --- | --- | --- | --- | --- |
| **Executive control** |  |  |  |  |  |
| Intercept | .219 | .005 | [.21, .23] | 47.72 | <.001 |
| Age | -.002 | .0008 | [-.004, -.0005] | -2.64 | <.01 |
| Age<sup>2</sup> | .0004 | 0.0002 | [.0001, .0007] | 2.39 | <.05 |
| **Orienting** |  |  |  |  |  |
| Intercept | .119 | .004 | [.113, .126] | 32.20 | <.001 |
| Age | .002 | .0006 | [.0004, 0.003] | 2.66 | <.01 |
| Sex | -.018 | .005 | [-.029, -.008] | -3.40 | <.001 |
| **Alerting** |  |  |  |  |  |
| Intercept | .096 | .003 | [.090, .102] | 32.12 | <.001 |
| Age | -.002 | .0007 | [-.003, -.0006] | -2.93 | <.01 |

Notes. S.E = Standard Error, CI = Confidence interval

**Figure 1.**
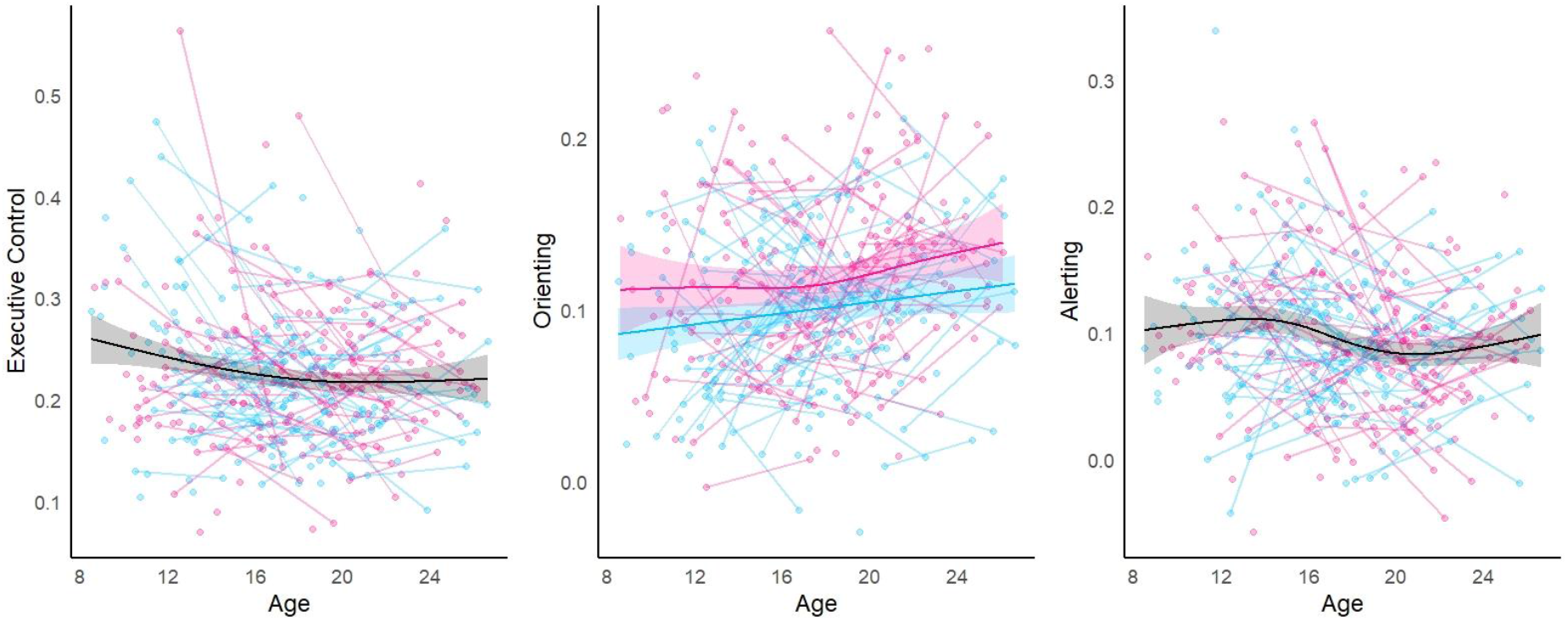
Developmental trajectories of the three attentional components. Left: a decelerating negative effect of age for executive control. Middle: a linear increase with age where females show higher scores in orienting than males. Right: a slight decrease in alerting during adolescence. Splines are plotted using generalized additive mixed modelling, and were plotted separately for each sex in orienting. Females are represented in pink and males in blue. Shaded areas correspond to the 95% confidence interval.

### Development of intraindividual variability of the ANT components

For executive control variability, none of the models were found to improve model fit compared to the null model (Table 7). Orienting variability (Table 8) and alerting variability (Table 9) were both best fitted by a linear age model with no main effect of sex or age-sex interaction effect. Thus, the results of the best fitting model for intraindividual variability scores in the ANT indicated development in two of the three attention networks (Table 10). Specifically, orienting variability was positively associated with age, while alerting variability was negatively associated with age. The GAMM slopes were in line with the statistical linear models and showed a linear increase and a linear decrease for orienting variability and alerting variability, respectively (Figure 2).

**Table 7.**
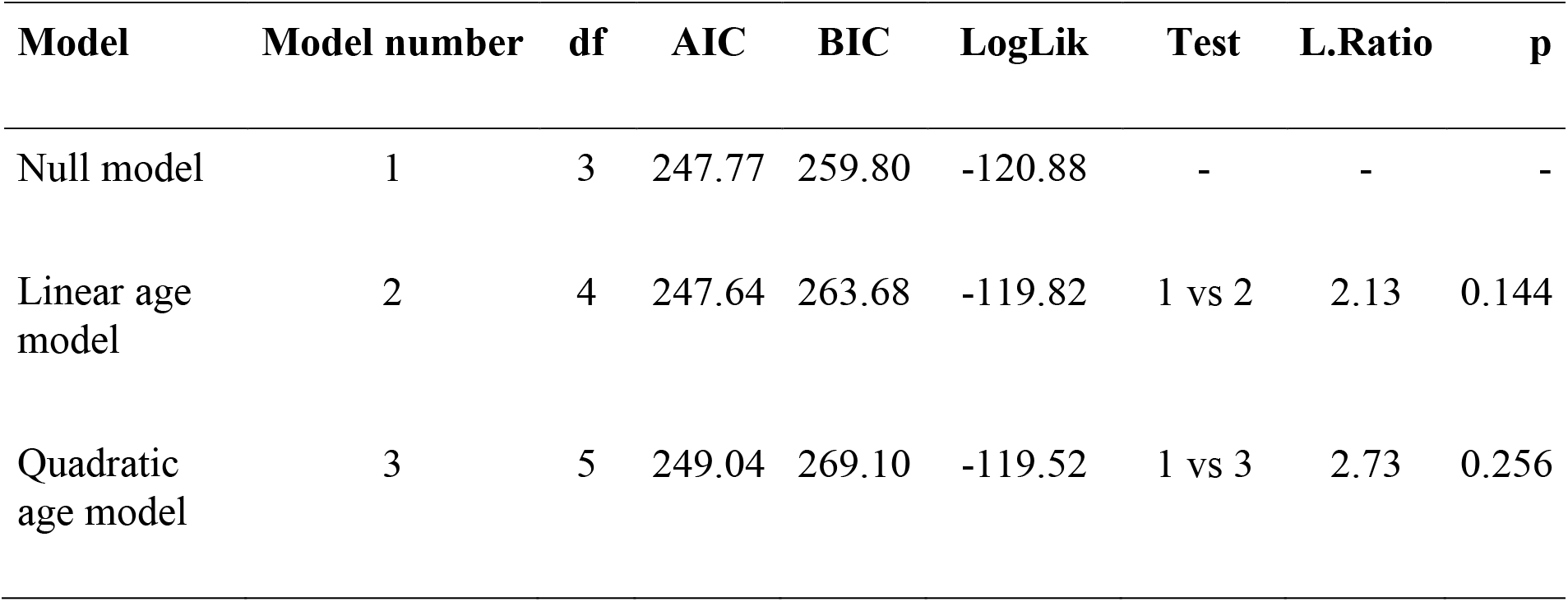
Comparison of models predicting executive control variability

**Table 8.** Comparison of models predicting orienting variability

| **Model** | **Model number** | **df** | **AIC** | **BIC** | **LogLik** | **Test** | **L.Ratio** | **p** |
| --- | --- | --- | --- | --- | --- | --- | --- | --- |
| Null model | 1 | 3 | 195.95 | 207.97 | -94.97 | - | - | - |
| Linear age model | 2 | 4 | 182.09 | 198.12 | -87.04 | 1 vs 2 | 15.86 | **<0.001** |
| Quadratic age model | 3 | 5 | 183.30 | 203.34 | -86.65 | 2 vs 3 | 0.78 | 0.376 |
| Main sex model | 4 | 5 | 183.91 | 203.96 | -86.96 | 2 vs 4 | 0.17 | 0.678 |
| Interaction sex*age model | 5 | 6 | 185.83 | 209.88 | -86.91 | 2 vs 5 | 0.26 | 0.880 |

**Table 9.** Comparison of models predicting alerting variability

| Model | Model number | df | AIC | BIC | LogLik | Test | L.Ratio | p |
| --- | --- | --- | --- | --- | --- | --- | --- | --- |
| Null model | 1 | 3 | 140.19 | 152.22 | -67.10 | - | - | - |
| Linear age model | 2 | 4 | 133.56 | 149.59 | -62.78 | 1 vs 2 | 8.63 | **0.003** |
| Quadratic age model | 3 | 5 | 135.53 | 155.57 | -62.76 | 2 vs 3 | 0.03 | 0.862 |
| Main sex model | 4 | 5 | 134.65 | 154.69 | -62.33 | 2 vs 4 | 0.91 | 0.341 |
| Interaction sex*age model | 5 | 6 | 136.24 | 160.30 | -62.12 | 2 vs 5 | 1.31 | 0.518 |

**Table 10.** Model parameters for the best fitting model of attention networks variability

| Attention network variability | Value | S.E | CI (95%)<br>[Lower, Upper] | t-value | p-value |
| --- | --- | --- | --- | --- | --- |
| **Orienting variability** |  |  |  |  |  |
| Intercept | .190 | .016 | [.160, .221] | 12.26 | <0.001 |
| Age | .015 | .004 | [.008, .022] | 4.02 | <.001 |
| **Alerting variability** |  |  |  |  |  |
| Intercept | .063 | .015 | [.035, .092] | 4.31 | <.001 |
| Age | -.010 | .003 | [-.017, -.003] | -2.95 | <0.01 |

Notes. S.E = Standard Error, CI = Confidence interval

**Figure 2.**
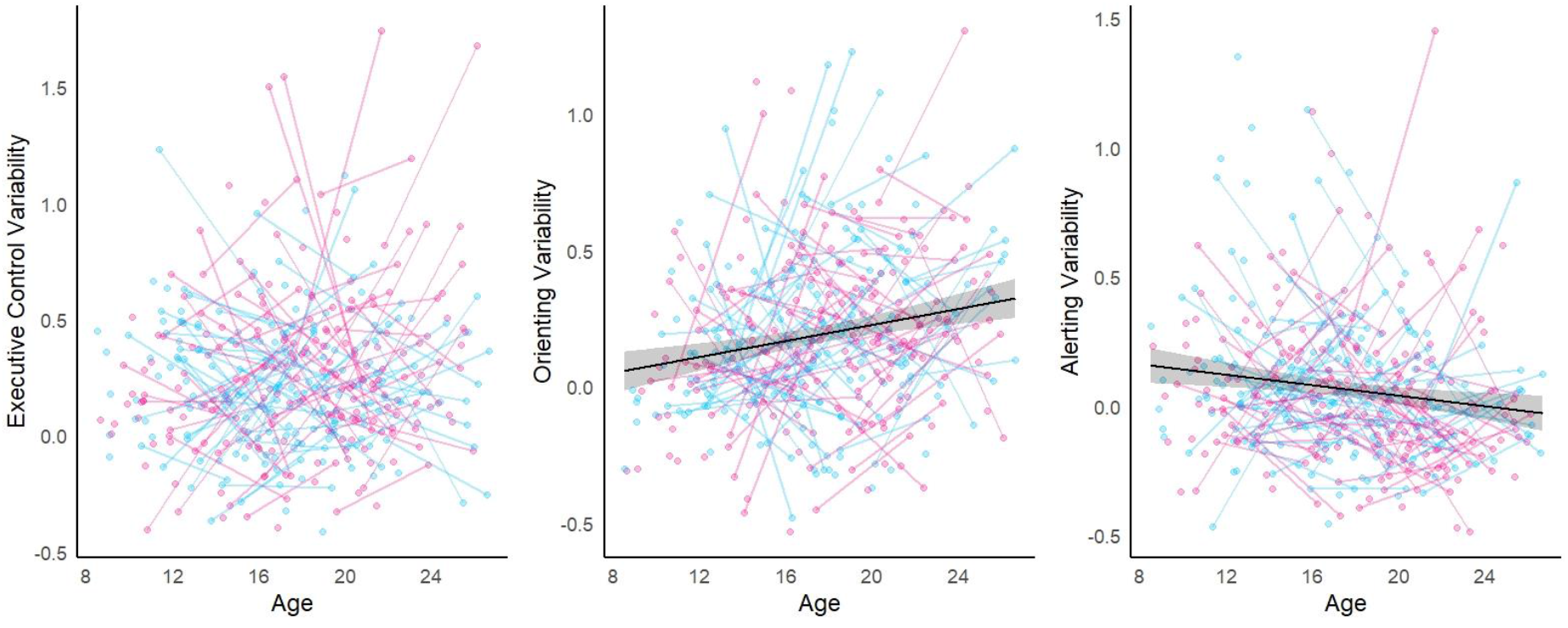
Developmental trajectories for the intraindividual variability measures of the three attentional components. Left: Spaghetti plot of executive control variability with no spline as no age terms improved the model fit of executive control variability. Middle: a linear increase with age on orienting variability. Right: a linear decrease with age on alerting variability. Splines are plotted using generalized additive mixed modelling. Shaded areas correspond to the 95% confidence interval.

### Associations between ANT performance and cortical thickness

There were significant regional associations between two out of six behavioral measures and cortical thickness (Figure 3). Higher alerting scores were associated with thicker cortex in a cluster encompassing parts of the right anterior prefrontal cortex, while higher orienting variability scores were associated with thinner cortex in two clusters including parts of the left anterior and superior prefrontal cortex. These main effects were largely the same without the age interaction terms included in the mixed effect model. Neither executive control or orienting, or executive control variability or alerting variability were associated with cortical thickness.

**Figure 3.**
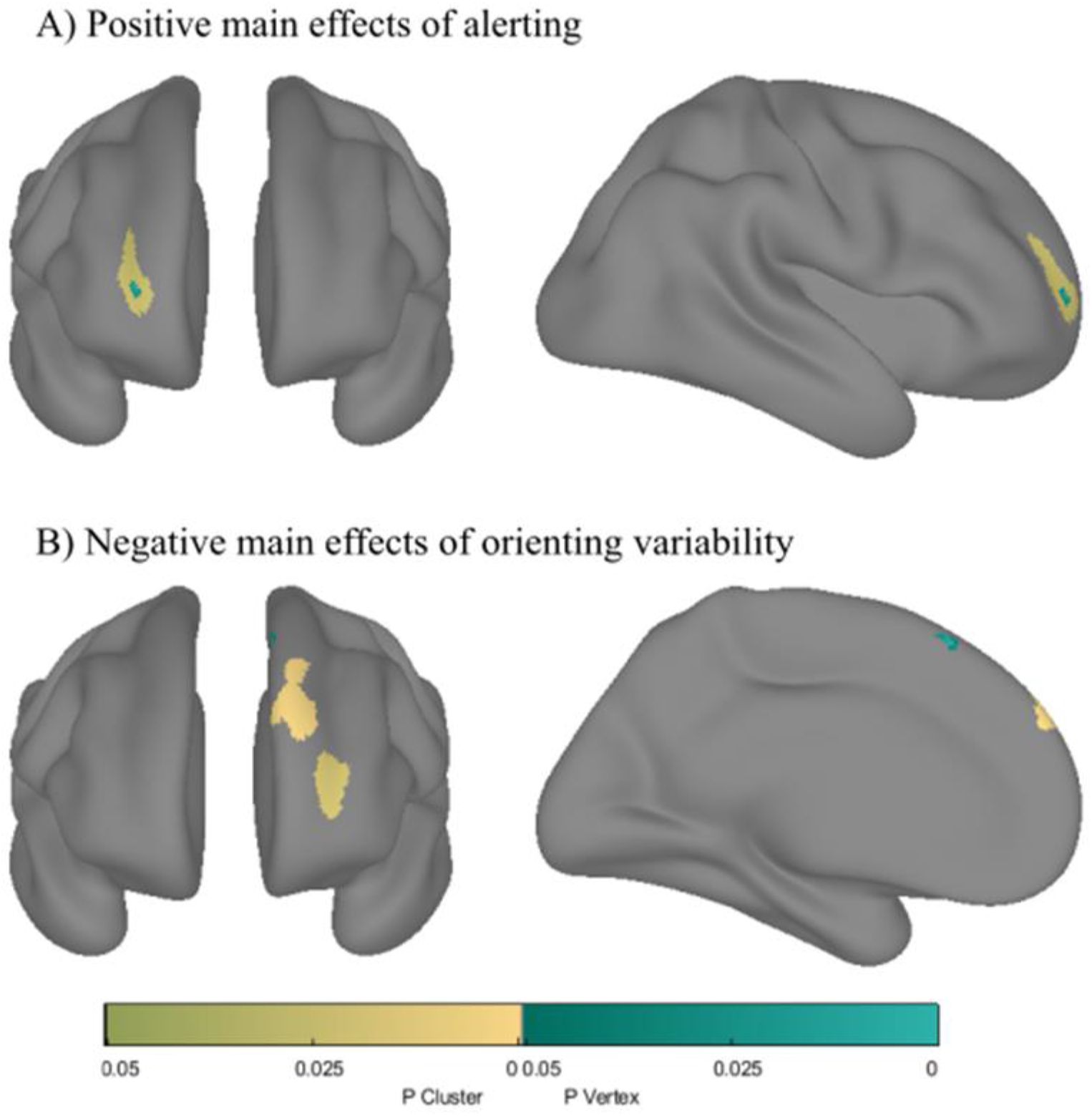
Main effects of the behavioral measures of the ANT on cortical thickness. A) A positive association between alerting and cortical thickness within the right anterior prefrontal cortex. B) A negative association between orienting variability and cortical thickness within the left anterior prefrontal cortex and left superior prefrontal cortex. The shades of yellow represent significant P-clusters, while the shades of turquoise represent significant vertices. The clusters and vertices are significant at p < .05 Random field theory corrected.

### Interactions between ANT performance and age on cortical thickness

The results of the analyses testing the interaction term between ANT performance and age was significant for three out of six of the behavioral measures (Figure 4). A negative interaction between orienting and age yielded three significant clusters within the left inferior parietal lobule, left dorsolateral prefrontal cortex and left insula. Furthermore, a negative interaction between alerting and age was associated with cortical thickness within the left rolandic operculum. A positive interaction between orienting variability and age yielded two significant clusters in the right insula and the right supplementary motor area. No significant age interactions with executive control, executive control variability, nor alerting variability were found on cortical thickness. Finally, no association was found between cortical thickness clusters and any of the ANT measures interacting with age^2^.

**Figure 4.**
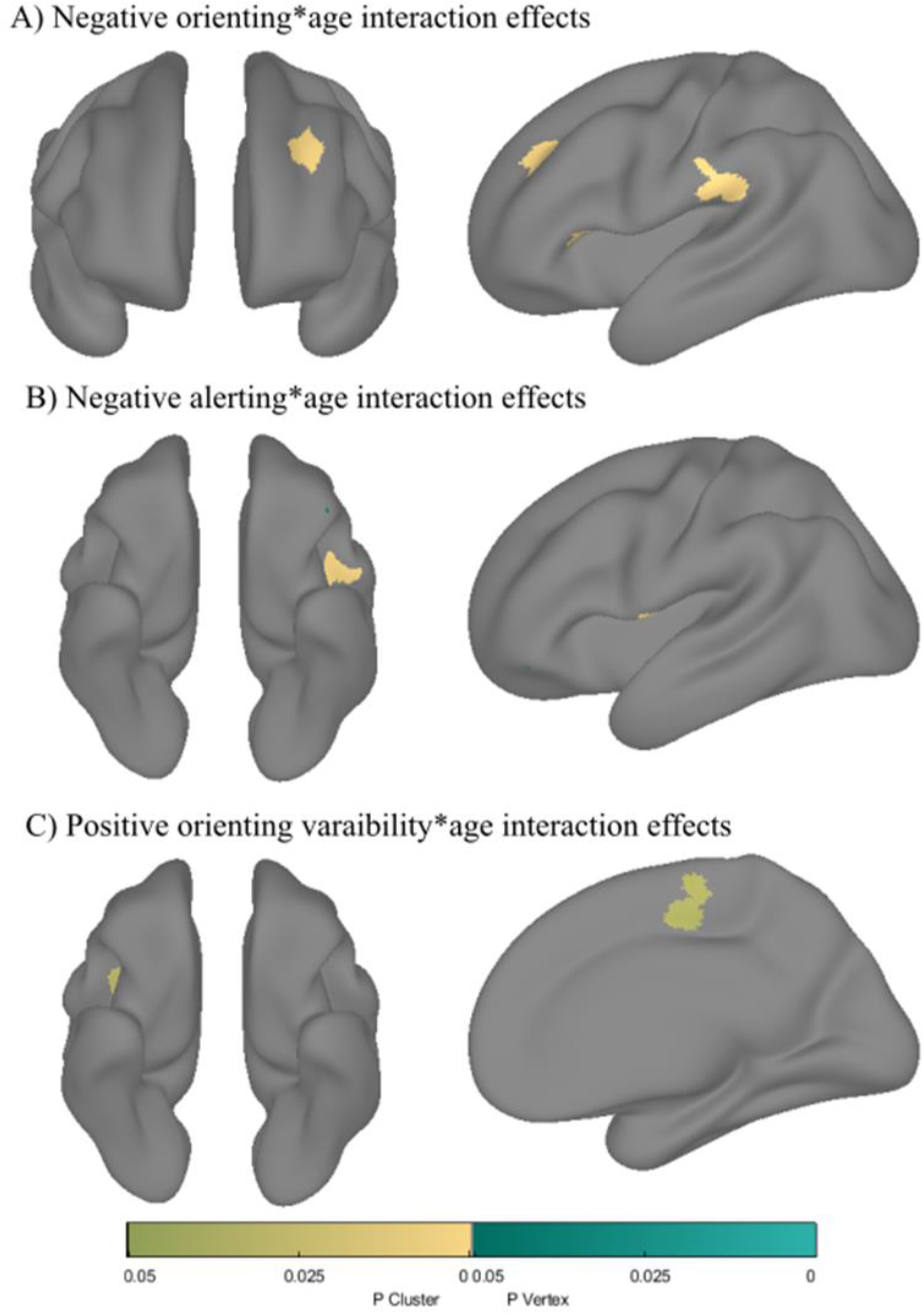
Interaction effect between the behavioral measures of the ANT and age on cortical thickness. A) Higher orienting scores were associated with greater age-related cortical thinning within the left dorsolateral prefrontal cortex, left insula and left inferior parietal lobule. B) Higher alerting scores were associated with greater age-related cortical thinning within left rolandic operculum. C) Lower orienting variability were associated with greater age-related cortical thinning within right insula and right medial supplementary motor area. The shades of yellow represent significant P-clusters, while the shades of turquoise represent significant vertices. The clusters and vertices are significant at p < .05 Random field theory corrected.

## Discussion

The present study investigated the developmental trajectories of three attentional functions and their intraindividual variability from late childhood to young adulthood, as well as their relationships to cortical thickness. Executive control showed a decelerating development from late childhood and into adolescence, with an initial improvement in late childhood and early adolescence, followed by a stabilization in late adolescence. Orienting continued to develop from late childhood to young adulthood, illustrated by a continuous increase in orienting performance, with females having higher orienting scores than males. Alerting did also exhibit a continuous development across the current age range, with a negative association with age. We also found a linear increase in orienting variability and a linear decrease in alerting variability with age. For the brain-behavior relationships, there was a main effect of two out of six of the behavioral measures on cortical thickness. Lower alerting scores were related to thinner cortex in a cluster within the right anterior prefrontal cortex, while higher orienting variability scores were related to thinner cortex in frontal regions. In addition, we identified an interaction effect with age on cortical thickness for three out of six of the behavioral measures. Here, higher orienting scores were related to greater age-related cortical thinning in the left inferior parietal and left prefrontal regions, higher alerting scores were related to greater age-related cortical thinning in the left operculum, and lower orienting variability scores were related to greater age-related cortical thinning in the right insula and the right motor area.

In line with our predictions and previous research that has shown age-related changes in executive control during adolescence (Waszak et al., 2010), our results support development of executive control that seem to stabilize around late adolescence. However, in contrast to previous research that has shown orienting to be fully developed by late childhood (Mullane et al., 2016; Pozuelos et al., 2014; Suades-González et al., 2017), we found evidence for further age-related changes in orienting, where higher orienting scores were associated with higher ages. Most notably, we identified a main effect of sex with females having higher orienting scores compared to males, which could indicate that females benefit more from spatial cues than males. Furthermore, the results indicated a linear decrease in alerting with age, although the GAMM showed a more complex trajectory as the youngest participants seemed to profit from the alerting cues, while its beneficial effect seemed to decelerate during adolescence. As previously suggested, this might be due to poor performance in no cue trials among the youngest participants. (Rueda et al., 2004). It should also be noted that previous research on development of attention networks using ANT have included younger samples and used a modified version of ANT, such as the child version (Mezzacappa, 2004; Rueda et al., 2004; Suades-González et al., 2017). Hence, the discrepancies in the two versions could explain inconsistencies between the developmental trajectories of the attention networks reported with the child version and our results from the adult version.

It is well known that intraindividual variability of the reaction times in cognitive tasks show large age-effects across childhood and adolescence (Dykiert et al., 2012; Tamnes et al., 2012). While there have been several studies investigating the development of the attentional components, studies mapping the developmental trajectory of the intraindividual variability in the attentional components are lacking. To fill this knowledge gap, we substituted the median reaction time with standard deviation when calculating attention network performance, hence calculating estimates for intraindividual variability in the three attentional components. Although we expected a decrease in executive control variability with age, the results did not indicate any age-related effects. However, in line with our predictions, there was a linear increase with age on orienting variability, indicating increased consistency in spatial cue trials with age. Further, the analyses also revealed that the alerting variability decreased with age and it seems plausible that this is due to faster increase in consistency in no cue trials, which would support the previous notion that youngest participants perform worse when there is no cues (Rueda et al., 2004), hence slower reaction times and larger trial-to-trial variability in no cue trials.

Intraindividual variability measures may reflect attentional lapses during cognitive tasks, which might be more common in children (Tamnes et al., 2012) and in individuals with ADHD (Adams et al., 2011). In fact, variability measures of reaction times often provide robust group differences between children with ADHD and control groups, irrespective of the cognitive task at hand, and have greater effect sizes compared to many other neuropsychological test outcomes (Epstein et al., 2011). Thus, we urge future studies to further utilize the present intraindividual variability measures obtained from the ANT as this could provide further insight into the development of performance consistency and differences in variability in participants with attentional problems.

We also investigated the relationship between behavioral measures obtained from the ANT and cortical thickness. First, we examined the main effects of the behavioral measures on cortical thickness. Although previous structural studies have found associations between executive control and anterior cingulate and the prefrontal cortex (Westlye et al., 2011), we did not observe any significant relationship between executive control and cortical thickness. However, in line with our prediction, we did observe a main effect of alerting in frontal regions, although no relations to thickness in parietal regions were found. Here, lower alerting scores were related to thinner cortex in a cluster within the right anterior prefrontal region. However, in contrast to our positive association between alerting and cortical thickness, Westlye and colleagues (2011) did find a negative association between cortical thickness in parietal regions and alerting among adults. To reconcile these results, it seems plausible that lower alerting scores are indeed indicative of better attentional functioning, while cortical thinning is part of typical development in youths (Tamnes et al., 2017) and reflects aging processes in older adults (Fjell et al., 2009), which could explain why there is a positive association between alerting and cortical thickness early in life, and a negative association between alerting and cortical thickness later in life. In addition, assuming that reduced cortical thickness reflects better cognitive functioning among youths, this would further support the negative association between orienting variability scores and cortical thickness in left prefrontal regions, as higher orienting variability scores indicate increased consistency in spatial cue trials compared to the center cue trials.

The results also yielded significant clusters from the age interaction term with orienting, alerting and the orienting variability. It is interesting to note that orienting showed a different developmental pattern than predicted. Firstly, we did not expect any age-related differences during adolescence in the behavioral performance, while we predicted orienting performance to be associated with cortical thickness in superior parietal regions. However, in addition to exhibit age-related improvements, orienting was also related to two cortical clusters within prefrontal regions and one in the inferior parietal region. It is further interesting to note that increased orienting scores indicates increased age-related cortical thinning in the left insula, while the complete opposite pattern was observed for orienting variability, where higher orienting variability scores indicated less age-related cortical thinning in the right insula. Although speculative, this could indicate a role of the bilateral insula in the detection of relevant stimuli and in a speed-consistency trade off. Future studies should aim to investigate the role of the bilateral insula in orientation and possible role in speed and consistency during development. Finally, several of the significant clusters across alerting, orienting and orienting variability performance showed interaction with age in predominantly frontal regions, which might reflect a general improvement of higher attentional functioning during maturation of the frontal cortex. However, as we do not know the direction of causality or the underlying mechanisms of cortical thinning, we urge caution when interpreting these results. Indeed, it is important to note that the age-related cortical thinning should not be interpreted as thinning per se, but as MRI apparent cortical thinning of the distance between the estimated outer grey and white matter surfaces (Walhovd et al., 2017). Apparent cortical thinning with increasing age during development can partly be explained by increased myelination of the inner layer of the grey matter with age (Natu et al., 2019). Increased myelination could result in more efficient neural processing, which is likely to be part of the underlying mechanism that allow for the observed continuous reduction in both reaction time and intraindividual variability during development.

To summarize, the current study included cross sectional and longitudinal data that provides important insight into the developmental trajectories of the efficacy of three attention networks. The behavioral results showed a decelerating age effect on executive control, stabilizing around late adolescence. Orienting showed continued development into young adulthood, with females having higher orienting scores than males, while alerting showed subtle, but more complex developmental patterns. Orienting variability showed a positive linear age-effect, while alerting variability exhibited a negative linear age-effect across late childhood and young adulthood. Further, lower alerting scores were associated with thinner cortex within a cluster encompassing parts of the right anterior prefrontal cortex, while higher orienting variability scores were associated with thinner cortex in clusters within the left anterior prefrontal and left superior prefrontal cortex. Finally, higher orienting scores were related to greater age-related cortical thinning in left dorsolateral prefrontal cortex, left insula and left inferior parietal lobule, higher alerting was related to greater age-related thinning in left rolandic operculum, while higher orienting variability scores were related to less age-related cortical thinning in the right insula and right medial supplementary motor area.

## Acknowledgements

This study was supported by the Research Council of Norway (#230345, #288083, #223273, # 186092, # 249931), Center for Lifespan Changes in Brain and Cognition, and the Norwegian Centre of Expertise for Neurodevelopmental Disorders and Hypersomnias.

## Supplementary materials

**Supplementary Figure 1.**
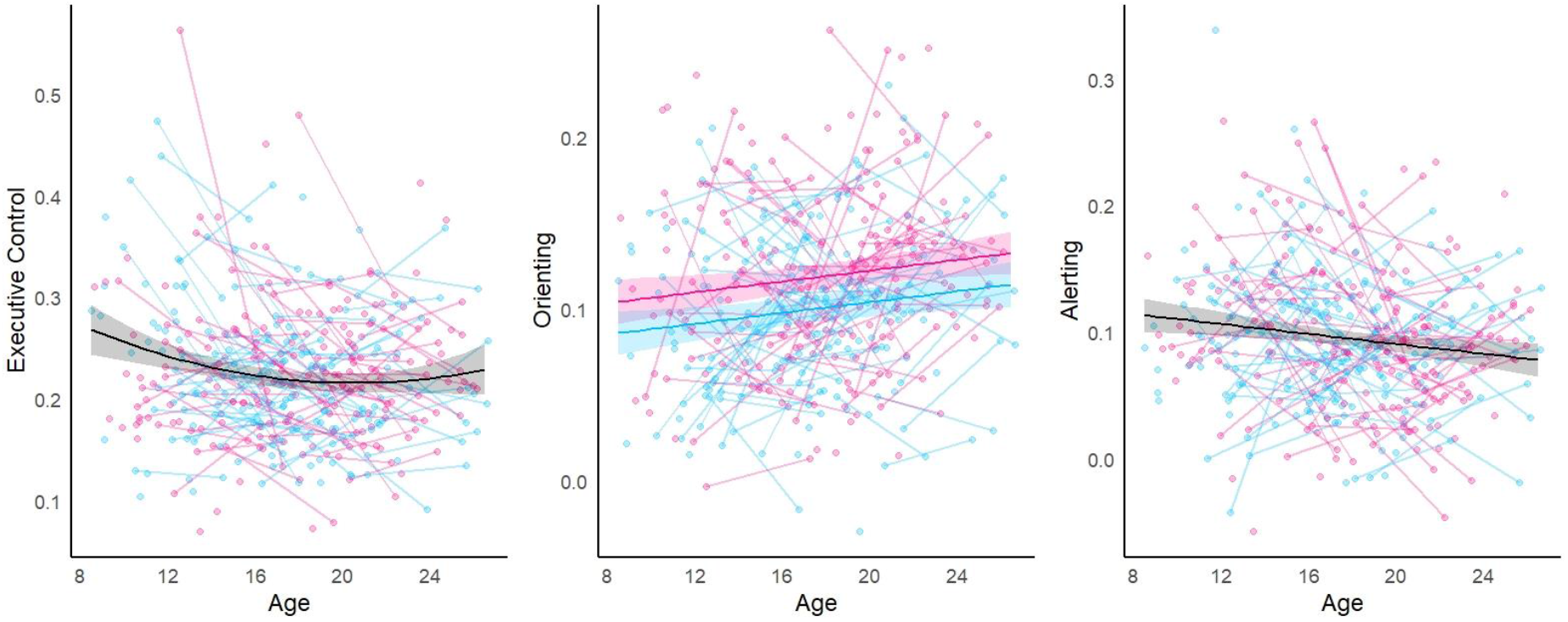
Developmental trajectories for the three attentional components. Left: a decelerating negative effect of age on executive control. Middle: a linear positive effect of age and a main effect of sex on orienting. Right: a negative linear effect of age on alerting. The regression lines are plotted using the mixed models that best fitted the data. Shaded areas correspond to the 95% confidence interval. Boys are represented in blue and girls are represented in pink.

**Supplementary Figure 2.**
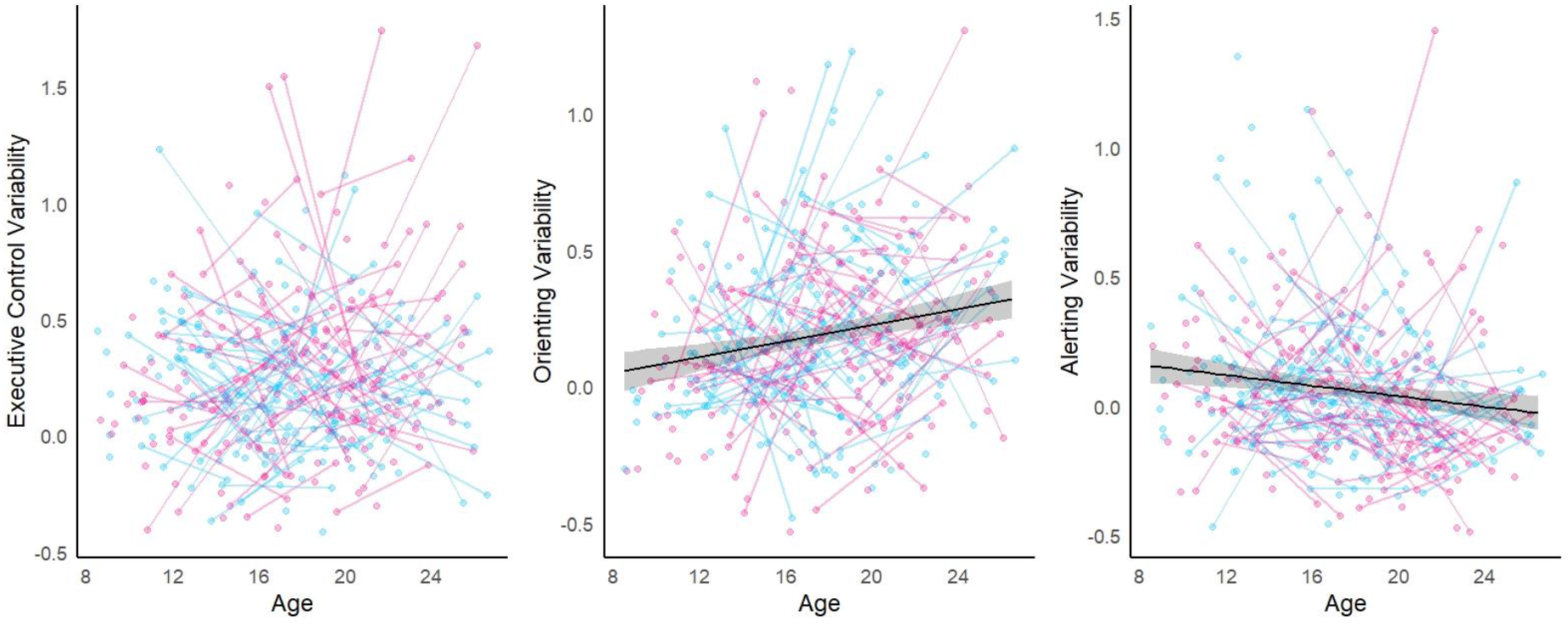
Developmental trajectories for the intraindividual variability measures of the three attentional components. Left: spaghetti plot of executive control variability with no regression line as no age terms improved the model fit of executive control variability. Middle, a positive linear effect of age on orienting variability. Right, a negative linear effect of age on alerting variability. Shaded areas correspond to the 95% confidence interval. The regression lines are plotted using the mixed models that best fitted the data.

## References

Adams, Z. W., Roberts, W. M., Milich, R., & Fillmore, M. T. (2011). Does response variability predict distractibility among adults with attention-deficit/hyperactivity disorder? Psychological Assessment, 23(2), 427–436. https://doi.org/10.1037/a0022112

Adólfsdóttir, S., Sørensen, L., & Lundervold, A. J. (2008). The attention network test: A characteristic pattern of deficits in children with ADHD. Behavioral and Brain Functions, 4(1), 9. https://doi.org/10.1186/1744-9081-4-9

Baijal, S., Jha, A. P., Kiyonaga, A., Singh, R., & Srinivasan, N. (2011). The Influence of Concentrative Meditation Training on the Development of Attention Networks during Early Adolescence. Frontiers in Psychology, 2. https://doi.org/10.3389/fpsyg.2011.00153

Bielak, A. A. M., Hultsch, D. F., Strauss, E., Macdonald, S. W. S., & Hunter, M. A. (2010). Intraindividual variability in reaction time predicts cognitive outcomes 5 years later. Neuropsychology, 24(6), 731–741. https://doi.org/10.1037/a0019802

Dale, A. M., Fischl, B., & Sereno, M. I. (1999). Cortical Surface-Based Analysis: I. Segmentation and Surface Reconstruction. NeuroImage, 9(2), 179–194. https://doi.org/10.1006/nimg.1998.0395

Dennis, E. L., & Thompson, P. M. (2013). Typical and atypical brain development: A review of neuroimaging studies. Dialogues in Clinical Neuroscience, 15(3), 359–384.

Dykiert, D., Der, G., Starr, J. M., & Deary, I. J. (2012). Sex differences in reaction time mean and intraindividual variability across the life span. Developmental Psychology, 48(5), 1262. https://doi.org/10.1037/a0027550

Epstein, J. N., Langberg, J. M., Rosen, P. J., Graham, A., Narad, M. E., Antonini, T. N., Brinkman, W. B., Froehlich, T., Simon, J. O., & Altaye, M. (2011). Evidence for higher reaction time variability for children with ADHD on a range of cognitive tasks including reward and event rate manipulations. Neuropsychology, 25(4), 427. https://doi.org/10.1037/a0022155

Eriksen, B. A., & Eriksen, C. W. (1974). Effects of noise letters upon the identification of a target letter in a nonsearch task. Perception & Psychophysics, 16(1), 143–149. https://doi.org/10.3758/BF03203267

Fan, J., Bernardi, S., Dam, N. T., Anagnostou, E., Gu, X., Martin, L., Park, Y., Liu, X., Kolevzon, A., Soorya, L., Grodberg, D., Hollander, E., & Hof, P. R. (2012). Functional deficits of the attentional networks in autism. Brain and Behavior, 2(5), 647–660. https://doi.org/10.1002/brb3.90

Fan, J., Gu, X., Guise, K. G., Liu, X., Fossella, J., Wang, H., & Posner, M. I. (2009). Testing the behavioral interaction and integration of attentional networks. Brain and Cognition, 70(2), 209–220. https://doi.org/10.1016/j.bandc.2009.02.002

Fan, J., McCandliss, B. D., Fossella, J., Flombaum, J. I., & Posner, M. I. (2005). The activation of attentional networks. NeuroImage, 26(2), 471–479. https://doi.org/10.1016/j.neuroimage.2005.02.004

Fan, J., McCandliss, B. D., Sommer, T., Raz, A., & Posner, M. I. (2002). Testing the Efficiency and Independence of Attentional Networks. Journal of Cognitive Neuroscience, 14(3), 340–347. https://doi.org/10.1162/089892902317361886

Federico, F., Marotta, A., Martella, D., & Casagrande, M. (2017). Development in attention functions and social processing: Evidence from the Attention Network Test. British Journal of Developmental Psychology, 35(2), 169–185. https://doi.org/10.1111/bjdp.12154

Ferschmann, L., Vijayakumar, N., Grydeland, H., Overbye, K., Sederevicius, D., Due-Tønnessen, P., Fjell, A. M., Walhovd, K. B., Pfeifer, J. H., & Tamnes, C. K. (2019). Prosocial behavior relates to the rate and timing of cortical thinning from adolescence to young adulthood. Developmental Cognitive Neuroscience, 40, 100734. https://doi.org/10.1016/j.dcn.2019.100734

Fischl, B., Sereno, M. I., & Dale, A. M. (1999). Cortical surface-based analysis. II: Inflation, flattening, and a surface-based coordinate system. NeuroImage, 9(2), 195–207. https://doi.org/10.1006/nimg.1998.0396

Fischl, Bruce. (2012). FreeSurfer. NeuroImage, 62(2), 774–781. https://doi.org/10.1016/j.neuroimage.2012.01.021

Fischl, Bruce, Salat, D. H., Busa, E., Albert, M., Dieterich, M., Haselgrove, C., van der Kouwe, A., Killiany, R., Kennedy, D., Klaveness, S., Montillo, A., Makris, N., Rosen, B., & Dale, A. M. (2002). Whole Brain Segmentation: Automated Labeling of Neuroanatomical Structures in the Human Brain. Neuron, 33(3), 341–355. https://doi.org/10.1016/S0896-6273(02)00569-X

Fjell, A. M., Grydeland, H., Krogsrud, S. K., Amlien, I., Rohani, D. A., Ferschmann, L., Storsve, A. B., Tamnes, C. K., Sala-Llonch, R., Due-Tønnessen, P., Bjørnerud, A., Sølsnes, A. E., Håberg, A. K., Skranes, J., Bartsch, H., Chen, C.-H., Thompson, W. K., Panizzon, M. S., Kremen, W. S., … Walhovd, K. B. (2015). Development and aging of cortical thickness correspond to genetic organization patterns. Proceedings of the National Academy of Sciences of the United States of America, 112(50), 15462–15467. https://doi.org/10.1073/pnas.1508831112

Fjell, A. M., Walhovd, K. B., Westlye, L. T., Østby, Y., Tamnes, C. K., Jernigan, T. L., Gamst, A., & Dale, A. M. (2010). When does brain aging accelerate? Dangers of quadratic fits in cross-sectional studies. NeuroImage, 50(4), 1376–1383. https://doi.org/10.1016/j.neuroimage.2010.01.061

Fjell, A. M., Westlye, L. T., Amlien, I., Espeseth, T., Reinvang, I., Raz, N., Agartz, I., Salat, D. H., Greve, D. N., Fischl, B., Dale, A. M., & Walhovd, K. B. (2009). High Consistency of Regional Cortical Thinning in Aging across Multiple Samples. Cerebral Cortex, 19(9), 2001–2012. https://doi.org/10.1093/cercor/bhn232

Galvao-Carmona, A., González-Rosa, J. J., Hidalgo-Muñoz, A. R., Páramo, D., Benítez, M. L., Izquierdo, G., & Vázquez-Marrufo, M. (2014). Disentangling the attention network test: Behavioral, event related potentials, and neural source analyses. Frontiers in Human Neuroscience, 8, 813. https://doi.org/10.3389/fnhum.2014.00813

Gopalan, P. R. S., Loberg, O., Hämäläinen, J. A., & Leppänen, P. H. T. (2019). Attentional processes in typically developing children as revealed using brain event-related potentials and their source localization in Attention Network Test. Scientific Reports, 9(1), 2940. https://doi.org/10.1038/s41598-018-36947-3

Herting, M. M., Johnson, C., Mills, K. L., Vijayakumar, N., Dennison, M., Liu, C., Goddings, A.-L., Dahl, R. E., Sowell, E. R., Whittle, S., Allen, N. B., & Tamnes, C. K. (2018). Development of subcortical volumes across adolescence in males and females: A multisample study of longitudinal changes. NeuroImage, 172, 194–205. https://doi.org/10.1016/j.neuroimage.2018.01.020

Houston, S. M., Herting, M. M., & Sowell, E. R. (2014). The neurobiology of childhood structural brain development: Conception through adulthood. Current Topics in Behavioral Neurosciences, 16, 3–17. https://doi.org/10.1007/7854_2013_265

Konrad, K., Neufang, S., Thiel, C. M., Specht, K., Hanisch, C., Fan, J., Herpertz-Dahlmann, B., & Fink, G. R. (2005). Development of attentional networks: An fMRI study with children and adults. NeuroImage, 28(2), 429–439. https://doi.org/10.1016/j.neuroimage.2005.06.065

Lewis, F. C., Reeve, R. A., & Johnson, K. A. (2016). A longitudinal analysis of the attention networks in 6-to 11-year-old children. Child Neuropsychology, 24(2), 145–165. https://doi.org/10.1080/09297049.2016.1235145

MacDonald, S. W. S., Li, S.-C., & Bäckman, L. (2009). Neural underpinnings of within-person variability in cognitive functioning. Psychology and Aging, 24(4), 792–808. https://doi.org/10.1037/a0017798

MacDonald, S. W. S., Nyberg, L., & Bäckman, L. (2006). Intra-individual variability in behavior: Links to brain structure, neurotransmission and neuronal activity. Trends in Neurosciences, 29(8), 474–480. https://doi.org/10.1016/j.tins.2006.06.011

Mahoney, J. R., Verghese, J., Goldin, Y., Lipton, R., & Holtzer, R. (2010). Alerting, orienting, and executive attention in older adults. Journal of the International Neuropsychological Society: JINS, 16(5), 877–889. https://doi.org/10.1017/S1355617710000767

Mezzacappa, E. (2004). Alerting, Orienting, and Executive Attention: Developmental Properties and Sociodemographic Correlates in an Epidemiological Sample of Young, Urban Children. Child Development, 75(5), 1373–1386. https://doi.org/10.1111/j.1467-8624.2004.00746.x

Mogg, K., Salum, G. A., Bradley, B. P., Gadelha, A., Pan, P., Alvarenga, P., Rohde, L. A., Pine, D. S., & Manfro, G. G. (2015). Attention network functioning in children with anxiety disorders, attention-deficit/hyperactivity disorder and non-clinical anxiety. Psychological Medicine, 45(12), 2633–2646. https://doi.org/10.1017/S0033291715000586

Mullane, J. C., Lawrence, M. A., Corkum, P. V., Klein, R. M., & McLaughlin, E. N. (2016). The development of and interaction among alerting, orienting, and executive attention in children. Child Neuropsychology, 22(2), 155–176. https://doi.org/10.1080/09297049.2014.981252

Natu, V. S., Gomez, J., Barnett, M., Jeska, B., Kirilina, E., Jaeger, C., Zhen, Z., Cox, S., Weiner, K. S., Weiskopf, N., & Grill-Spector, K. (2019). Apparent thinning of human visual cortex during childhood is associated with myelination. Proceedings of the National Academy of Sciences, 201904931. https://doi.org/10.1073/pnas.1904931116

Neuhaus, A. H., Urbanek, C., Opgen-Rhein, C., Hahn, E., Ta, T. M. T., Koehler, S., Gross, M., & Dettling, M. (2010). Event-related potentials associated with Attention Network Test. International Journal of Psychophysiology: Official Journal of the International Organization of Psychophysiology, 76(2), 72–79. https://doi.org/10.1016/j.ijpsycho.2010.02.005

Posner, M. I. (1980). Orienting of Attention. Quarterly Journal of Experimental Psychology, 32(1), 3–25. https://doi.org/10.1080/00335558008248231

Posner, M. I. (2008). Measuring Alertness. Annals of the New York Academy of Sciences, 1129(1), 193–199. https://doi.org/10.1196/annals.1417.011

Pozuelos, J. P., Paz-Alonso, P. M., Castillo, A., Fuentes, L. J., & Rueda, M. R. (2014). Development of attention networks and their interactions in childhood. Developmental Psychology, 50(10), 2405. https://doi.org/10.1037/a0037469

Reuter, M., Schmansky, N. J., Rosas, H. D., & Fischl, B. (2012). Within-subject template estimation for unbiased longitudinal image analysis. NeuroImage, 61(4), 1402–1418. https://doi.org/10.1016/j.neuroimage.2012.02.084

Rueda, M. R., Fan, J., McCandliss, B. D., Halparin, J. D., Gruber, D. B., Lercari, L. P., & Posner, M. I. (2004). Development of attentional networks in childhood. Neuropsychologia, 42(8), 1029–1040. https://doi.org/10.1016/j.neuropsychologia.2003.12.012

Spagna, A., Dong, Y., Mackie, M.-A., Li, M., Harvey, P. D., Tian, Y., Wang, K., & Fan, J. (2015). Clozapine improves the orienting of attention in schizophrenia. Schizophrenia Research, 168(1–2), 285–291. https://doi.org/10.1016/j.schres.2015.08.009

Suades-González, E., Forns, J., García-Esteban, R., López-Vicente, M., Esnaola, M., Álvarez-Pedrerol, M., Julvez, J., Cáceres, A., Basagaña, X., López-Sala, A., & Sunyer, J. (2017). A Longitudinal Study on Attention Development in Primary School Children with and without Teacher-Reported Symptoms of ADHD. Frontiers in Psychology, 8. https://doi.org/10.3389/fpsyg.2017.00655

Tamnes, C. K., Fjell, A. M., Westlye, L. T., Østby, Y., & Walhovd, K. B. (2012). Becoming Consistent: Developmental Reductions in Intraindividual Variability in Reaction Time Are Related to White Matter Integrity. Journal of Neuroscience, 32(3), 972–982. https://doi.org/10.1523/JNEUROSCI.4779-11.2012

Tamnes, C. K., Herting, M. M., Goddings, A.-L., Meuwese, R., Blakemore, S.-J., Dahl, R. E., Güroğlu, B., Raznahan, A., Sowell, E. R., Crone, E. A., & Mills, K. L. (2017). Development of the Cerebral Cortex across Adolescence: A Multisample Study of Inter-Related Longitudinal Changes in Cortical Volume, Surface Area, and Thickness. Journal of Neuroscience, 37(12), 3402–3412. https://doi.org/10.1523/JNEUROSCI.3302-16.2017

Tamnes, C.K., & Mills, K.L. (2020). Imaging structural brain development in childhood and adolescence. In R.M. George, D. Poeppel., & M.S. Gazzaniga (Eds.), The Cognitive Neurosciences (6th ed., pp. 17–25). MIT Press.

Tamnes, C. K., Østby, Y., Fjell, A. M., Westlye, L. T., Due-Tønnessen, P., & Walhovd, K. B. (2010). Brain Maturation in Adolescence and Young Adulthood: Regional Age-Related Changes in Cortical Thickness and White Matter Volume and Microstructure. Cerebral Cortex, 20(3), 534–548. https://doi.org/10.1093/cercor/bhp118

Tamnes, C. K., Walhovd, K. B., Dale, A. M., Østby, Y., Grydeland, H., Richardson, G., Westlye, L. T., Roddey, J. C., Hagler, D. J., Due-Tønnessen, P., Holland, D., & Fjell, A. M. (2013). Brain development and aging: Overlapping and unique patterns of change. NeuroImage, 68, 63–74. https://doi.org/10.1016/j.neuroimage.2012.11.039

Vijayakumar, N., Allen, N. B., Youssef, G., Dennison, M., Yücel, M., Simmons, J. G., & Whittle, S. (2016). Brain development during adolescence: A mixed-longitudinal investigation of cortical thickness, surface area, and volume. Human Brain Mapping, 37(6), 2027–2038. https://doi.org/10.1002/hbm.23154

Walhovd, K. B., Fjell, A. M., Giedd, J., Dale, A. M., & Brown, T. T. (2017). Through Thick and Thin: A Need to Reconcile Contradictory Results on Trajectories in Human Cortical Development. Cerebral Cortex, 27(2). https://doi.org/10.1093/cercor/bhv301

Waszak, F., Li, S.-C., & Hommel, B. (2010). The development of attentional networks: Cross-sectional findings from a life span sample. Developmental Psychology, 46(2), 337–349. https://doi.org/10.1037/a0018541

Westlye, L. T., Grydeland, H., Walhovd, K. B., & Fjell, A. M. (2011). Associations between Regional Cortical Thickness and Attentional Networks as Measured by the Attention Network Test. Cerebral Cortex, 21(2), 345–356. https://doi.org/10.1093/cercor/bhq101

Wierenga, L. M., Langen, M., Oranje, B., & Durston, S. (2014). Unique developmental trajectories of cortical thickness and surface area. NeuroImage, 87, 120–126. https://doi.org/10.1016/j.neuroimage.2013.11.010

Williams, B. R., Hultsch, D. F., Strauss, E. H., Hunter, M. A., & Tannock, R. (2005). Inconsistency in reaction time across the life span. Neuropsychology, 19(1), 88–96. https://doi.org/10.1037/0894-4105.19.1.88

Wood, S. (2017). Generalized additive models: An introduction with R. Chapman and Hall/CRC.

